# NMR structure verifies the eponymous degenerate zinc finger domain of transcription factor ZNF750

**DOI:** 10.1101/2023.06.10.544471

**Authors:** Antonio J. Rua, Richard D. Whitehead, Andrei T. Alexandrescu

## Abstract

ZNF750 is a nuclear transcription factor that activates skin differentiation and has tumor suppressor roles in several cancers. Unusually, ZNF750 has only a single zinc-finger (ZNF) domain, Z*, with an amino acid sequence that differs markedly from the CCHH family consensus. Because of its sequence differences Z* is classified as degenerate, presumed to have lost the ability to bind the zinc ion required for folding. AlphaFold predicts an irregular structure for Z* with low confidence. Low confidence predictions are often inferred to be intrinsically disordered regions of proteins, which would be the case if Z* did not bind Zn^2+^. We use NMR and CD spectroscopy to show that a 25-51 segment of ZNF750, corresponding to the Z* domain, folds into a well-defined antiparallel ββα tertiary structure with a pM dissociation constant for Zn^2+^, and a thermal stability >80 °C. Of three alternative Zn^2+^ ligand sets, Z* uses a CCHC rather than the expected CCHH motif. The switch in the last ligand maintains the folding topology and hydrophobic core of the classical ZNF motif. CCHC ZNFs are associated with protein-protein interactions but Z* binds DNA. Since the metal chelating site is on the other side of the molecule, it suggests functional preferences are a result of divergent evolution rather than physical constraints on the structure. The structure of Z* provides a context for understanding the domain’s DNA-binding properties and mutations associated with cancers. We expect other ZNFs currently classified as degenerate, are CCHC-type structures like Z*.

---

Transcription factors are sequence-specific DNA-binding proteins that control the transformation of genetic information from DNA to RNA. They account for ∼8% of human genes (1) and are extremely important in biology and health since they determine what, when, where, how much, and for how long genes are expressed. Nearly half of human transcription factors have small zinc-finger (ZNF) domains, typically in multiple copies (2). The ZNF domains usually adopt an antiparallel ββα folding motif upon binding divalent zinc (Zn^2+^). In transcription factors, ZNFs are used as base-pair recognition modules that enable site-specific binding to double-stranded DNA (dsDNA).

ZNF750, is a 723 a.a. nuclear transcription factor that mediates skin differentiation (3-6). Altered epidermal differentiation is a feature of more than 100 skin diseases, although the etiology is poorly understood at a mechanistic and molecular level (5,7,8). The ZNF750 gene was first discovered due to an autosomal dominant frameshift mutation within its only putative CCHH ZNF domain (9), that we will henceforth call Z*. The mutation occurred in a family that presented with seborrhea-like dermatosis with non-arthritic psoriasiform elements (SLDP), suggesting ZNF750 might play a role in genetic skin disease (9). This was subsequently confirmed by mutations in the promoter for ZNF750 that cause familial psoriasis (10,11). Recent work has shed light on the function of the transcription factor. ZNF750 is expressed in keratinocytes but not fibroblasts (5,9). Based on mass spectrometry and ChIP-seq data, ZNF750 is thought to have dual roles in epithelial homeostasis and skin differentiation (3-5). In concert with the chromatin regulator KDM1A, the ZNF750 transcription factor inhibits progenitor genes that regulate the proliferation of self-renewing keratinocytes. With the pluripotency transcription factor KLF4 (Krüppel-like factor 4), ZNF750 activates genes that control skin differentiation. The keratinocyte-differentiation function of ZNF750 has recently been shown to be involved in the epidermal inflammatory response, through its interaction with the S/T-kinase IRAK2 in psoriatic epidermis but not in healthy skin. Thus, ZNF750 could affect the severity of skin diseases (12).

Over the last few years it has become evident that in addition to skin differentiation ZNF750 has roles in oral (13) and esophageal (14) squamous cell carcinoma, melanoma (15), ocular sebaceous carcinoma (16), prostate (17) and breast cancers (18-20). A common mechanistic theme linking the involvement of ZNF750 in pathologies as varied as skin disease and cancers is that the transcription factor regulates cell differentiation. ZNF750 appears to act as a tumor suppressor by inhibiting cancer stem cells, and has additional roles in regulating tumor growth, cell migration, and adhesion (13). When ZNF750 is mutated or expressed at low levels, it no longer acts as a tumor suppressor and cancer cells proliferate (21).

Classical biological approaches have shed considerable light on the function of ZNF750. However there remain important open questions about the transcription factor that lend themselves to structural biology. ZNF750 has a single putative CCHH zinc-finger domain Z* (Fig. 1A), from which the transcription factor derives its name (5). The CCHH designation (sometimes also called C2H2) gives the order of the Cys, Cys, His, His Zn^2+^-ligands in the amino acid sequence. The vast majority of transcription factors have at least two or more CCHH ZNF domains, since a single copy is typically insufficient for sequence-specific DNA binding (22-24). Another unusual feature of the Z* domain is that it has an amino acid sequence that differs considerably from the ZNF consensus.

**Figure 1.**
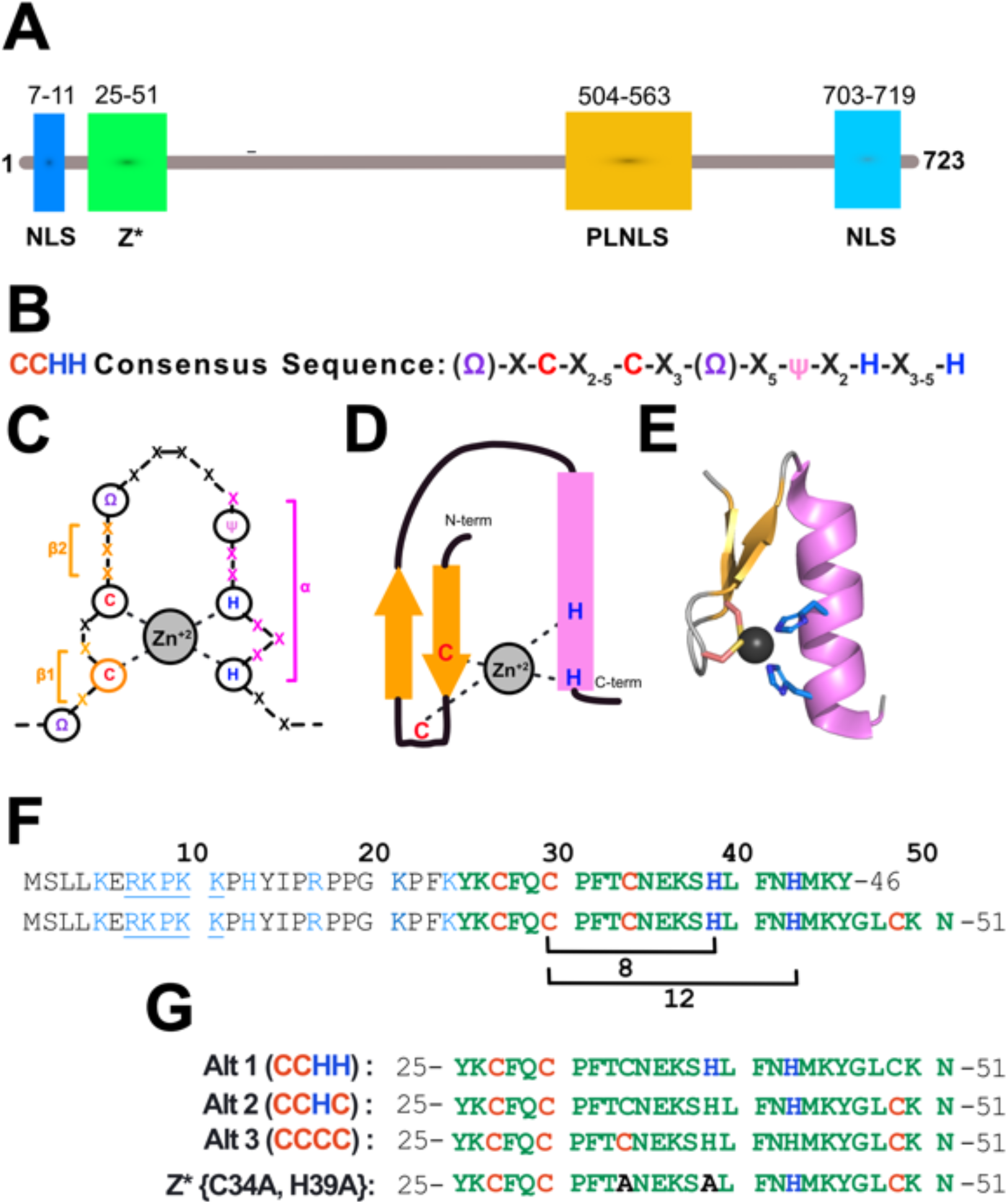
Sequence properties of ZNF750 and its Z* domain. **(A)** Domain diagram for ZNF750 adapted from (17). NLS, nuclear localization signal; PLNLS, sequence domains consisting of the given amino acids, thought to be important in mediating protein-protein interactions for ZNF750 (3); Z*, the putative ZNF domain that is the subject of this study. **(B)** The consensus sequence of a CCHH ZNF determines its Zn^2+^ coordination (**C**), secondary structure (**D**), and ββα folding motif structure (**E**). In B and C, Ω represents the aromatic residues Y or F, ψ hydrophobic residues, and X is any residue. (**F**) Sequence of the N-terminus of ZNF750. The first 30 residues have a high proportion of basic amino acids (light blue) some of which correspond to an NLS sequence (underlined). The sequence of the putative ZNF domain, Z*, is shown in green. The UniProt database gives the Z* domain boundary as 25-46. This leads to a highly unfavorable 8-residue spacing between the second and third Zn^2+^-ligands. Extending the domain boundary by five amino acids to residue 51 introduces an additional Cys. With the extended domain boundary, there are three alternative Zn^2+^-ligand sets (**G**). The second alternative CCHC, gives a more favorable 12-residue spacing between the second and third ligands. The Z*{C34A, H39A) mutant peptide was used to confirm the Zn^2+^-ligands identified from the NMR structure.

The presence of Cys and His residues that can chelate Zn^2+^ is not sufficient to specify a ZNF domain. ZNF750 has 15 Cys and 22 His residues. Clearly, the protein would not function properly if all the potential ligands bound Zn^2+^ non-selectively. Rather, ZNF domains have amino acid preferences beyond the requirement of Cys/His ligands to bind Zn^2+^ (25,26), that vary for different ZNF families described by the order of metal-chelating ligands, such as CCCC, CCHC, CCCH. These families are associated with distinct functions such as DNA, RNA, protein, or lipid binding (26-30). The consensus sequence for the classical DNA-binding CCHH ZNFs of transcription factors is (Ω)-X-C-X_2-5_-C-X_3_-(Ω)-X_5_-ψ-X_2_-H-X_3-5_-H (Fig. 1B). Here, Ω is the aromatic residues Y or F, ψ is a hydrophobic residue, X is any residue, and the subscripts indicate the spacing between residues (26,27). The conserved hydrophobic and aromatic residues (shown in pink and purple in Fig. 1B-C) are required to form a small hydrophobic core (25). The spacing between metal ligands is important in accommodating the antiparallel ββα secondary structure motif, as well as for satisfying functional requirements (Fig. 1D-E). The “finger region” in CCHH ZNFs refers to the 12-residue segment between the second Cys and the first His ligands of Zn^2+^ (Fig. 1C). This region contains the residues that encode the dsDNA-binding specificity of ZNFs (31).

Mutations altering the spacing between the Zn^2+^-chelating residues in CCHH-type ZNFs, or replacing conserved aromatic/hydrophobic residues, can destabilize the structure or abrogate Zn^2+^-binding (32,33). The UniProt database (34) designates the domain boundaries for Z*, the putative CCHH-type ZNF in ZNF750, as residues 25-46 (Fig. 1F). With this domain assignment, the ‘finger region’ of Z* would have an 8 rather than the consensus 12-residues spacing, and none of the aromatic or hydrophobic residues that typically stabilize the ββα-fold. Due to the differences from consensus the UniProt database classifies Z* as ‘degenerate’, signifying the domain has probably lost its ability to bind Zn^2+^ (34,35). The premiere method for protein structure prediction AlphaFold (36,37), only models about 12 α-helical residues of the 723 a.a. protein with high confidence and predicts an irregular structure for the Z* domain with low confidence (PDB code AF_AFQ32MQ0F1). Clearly the folding state of Z*, whether the domain is a genuine ZNF, or an intrinsically disordered region without the ability to bind Zn^2+^, is critical for understanding the structure and function of ZNF750 – a transcription factor that derives its name from its sole ZNF domain. We therefore investigated the structure of Z* by nuclear magnetic resonance (NMR).

## RESULTS

### The Z* domain folds in the presence of Zn^2+^

Anticipating the Z* domain may not be a CCHH-type ZNF because of the resulting unfavorable 8-residue spacing between the second and third Zn^2+^-ligands, we synthesized a slightly longer fragment of ZNF750, from residues 25-51 (Fig. 1F). The longer fragment included C49, which could give a conventional 12-residue finger spacing if the second and third Zn^2+^ ligands were C30 and H43 (Fig. 1F). However, this would result in a CCHC rather than a CCHH set of Zn^2+^-ligands. A third possibility with the inclusion of C49 is a CCCC-type ZNF, since these are generally the least constrained in terms of sequence and ligand-spacing requirements. The three alternative types of Zn^2+^-ligand sets for the 25-51 fragment of ZNF750 are given in Fig. 1G.

Circular dichroism (CD) and NMR spectra of the synthetic 25-51 fragment of ZNF750 showed clear evidence of protein folding in the presence of a 1.2 molar excess of Zn^2+^ (Fig. 2).

**Figure 2.**
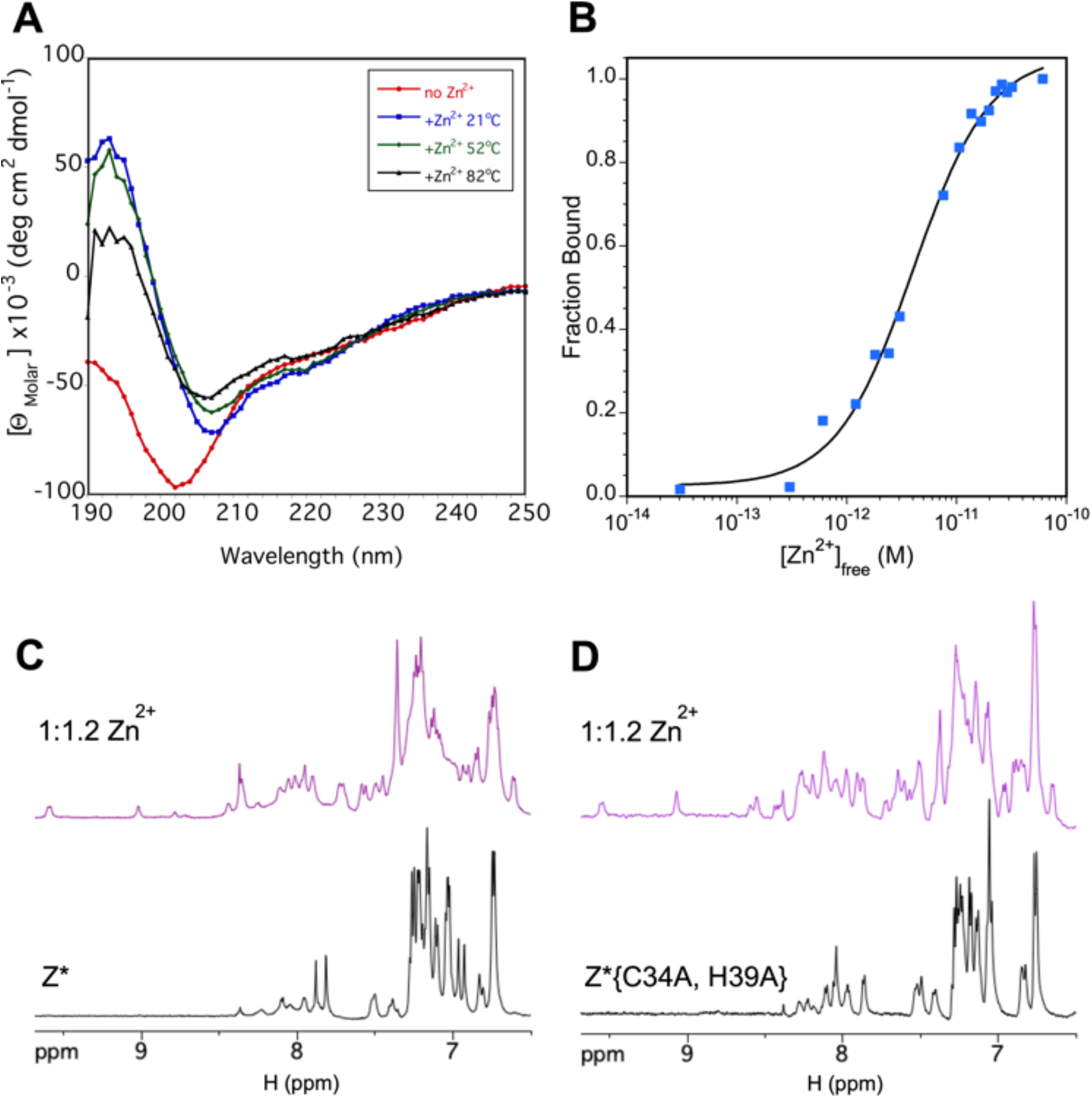
The Z* domain binds Zn^2+^ to form a stable folded structure. **(A)** CD spectra in the absence of Zn^2+^ (red) and in the presence of the metal at different temperatures. Note that the CD spectrum does not revert to that of the unfolded domain even at 82 °C. (**B**) Zn^2+^ binding curve for Z* in the presence of 22 mM EGTA. The data were fit to a sigmoidal curve, giving a K_d_ of 4.0 ± 0.4 x 10^−12^ M, and a Hill coefficient of 1.2 ± 0.1. (**C**) Downfield region of the ^1^H-NMR spectrum of Z* in the absence (black) and presence of Zn^2+^ (purple). (**D**) Same as C except with the C34A, H39A mutant of Z* used to verify the Zn^2+^-chelating residues determined from the NMR structure.

The CD spectrum is typical of an unfolded polypeptide in the absence of the metal but when Zn^2+^ is added matches that of folded ZNF domains (38,39) with a mixture of α-helix and β-sheet secondary structure (Fig. 2A). Consistently, addition of Zn^2+^ leads to increased chemical shift dispersion in the NMR spectrum, and enhanced amide proton signals due to reduced solvent exchange, indicating the domain adopts a stable tertiary structure (Fig. 2C).

To assess the stability of the Zn^2+^-bound Z* domain at pH 7, we varied the temperature from 21 to 82 °C in 4-degree increments. For clarity, data are only shown at three representative temperatures in Fig. 2A. With increasing temperature, the CD spectra show a linear increase in ellipticity at 208 nm and a decrease at 195 nm, that likely represents a pre-denaturation transition, rather than the sigmoidal transition typical of protein unfolding. The overall CD spectrum remains characteristic of a folded polypeptide up to 82 °C, and very different from that of the unfolded domain in the absence of Zn^2+^. We repeated the temperature experiment at pH 5.8, reasoning more acidic conditions could destabilize metal binding. We obtained similar results, however, with Z* remaining folded up to 90 °C. We conclude that the Zn^2+^-bound Z* domain is highly stable to thermal unfolding.

To assess metal binding affinity we carried out a competition experiment with the metal chelator EGTA (40). The use of the competing chelator was necessary because Z* binds Zn^2+^ too strongly to measure affinity directly. The binding data gives a K_d_ of 4 x 10^−12^ M for Zn^+2^ at pH 7.0 (Fig 2B). This value is within the range for ZNFs, that typically have K_d_ < 10^−9^ M but in some cases as small as 10^−18^ M (41-43).

In summary, the present results show that Z* is a genuine rather than degenerate ZNF. Upon binding Zn^2+^ with a pM affinity, the domain folds into a well-defined tertiary structure, with a thermal stability greater than 80-90 °C.

### Z* has a typical ZNF ββα fold

The Z* domain in the presence of Zn^2+^ gives excellent NMR data (Fig. 3). Each amino acid gives a single set of NMR resonances unlike other ZNF domains that exhibit spectral heterogeneity near histidines, due to interchange between aromatic ring Nε² and Nε¹ atom coordination of Zn^2+^ (44,45). We were able to obtain virtually complete NMR assignments of Z* using standard 2D TOCSY, NOESY (Fig. 3A), DQF-COSY and E-COSY experiments, together with natural abundance ^13^C-HSQC (Fig. 3B) and ^15^N-sofast HMQC (Fig. 3C) spectra. The NMR chemical shifts, together with short-range backbone NOE patterns, enabled us to calculate (46,47) the secondary structure of Z* shown in Figure 4. The calculated ββα secondary structure is a typical for a ZNF domain (48). We used the Talos-N program to predict S^2^ order parameters from the chemical shifts of Z*. The S^2^ order parameter ranges from 1 for rigid sites to 0 for sites with unrestricted flexibility. Except for the last two residues in the sequence K50 and N51, all of the S^2^ values for Z* domain were above 0.7. The muted flexibility of Z* is consistent with its high thermal stability.

**Figure 3.**
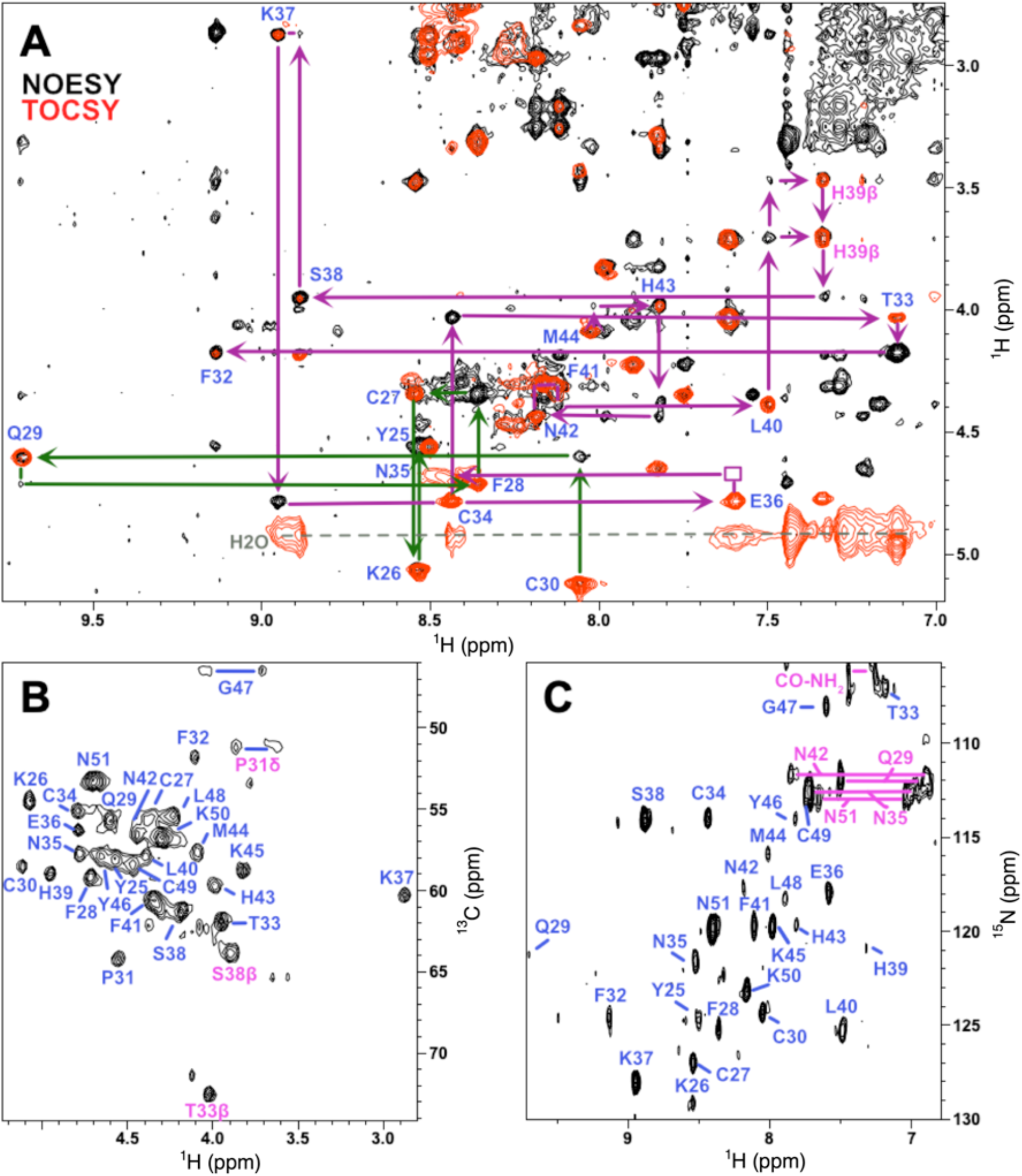
Representative 2D-NMR data for the Z* domain. **(A)** Superposition of TOCSY (red) and NOESY (black) spectra, illustrating sequential walks between residues M44 to F32 (purple) and C30 to Y25 (green) used to obtain NMR assignments. Blue labels indicate TOCSY H_N_-Hα crosspeaks, pink labels are sidechain crosspeaks. The group of peaks connected by a dashed gray line marked H_2_O at 4.93 ppm, are due rapid magnetization exchange between amide protons and water during the 70 ms TOCSY mixing time. (**B**) Fingerprint region of a natural-abundance ^1^H-^13^C HSQC spectrum. Blue labels indicate ^1^Hα-^13^Cα crosspeaks. (**C**) Natural-abundance ^1^H-^15^N sofast-HMQC spectrum. Blue labels indicate backbone ^1^H-^15^N correlations. Pink labels indicate correlations from sidechain -NH_2_ groups, and from the backbone C-terminal amide blocking group (CO-NH_2_).

**Figure 4.**
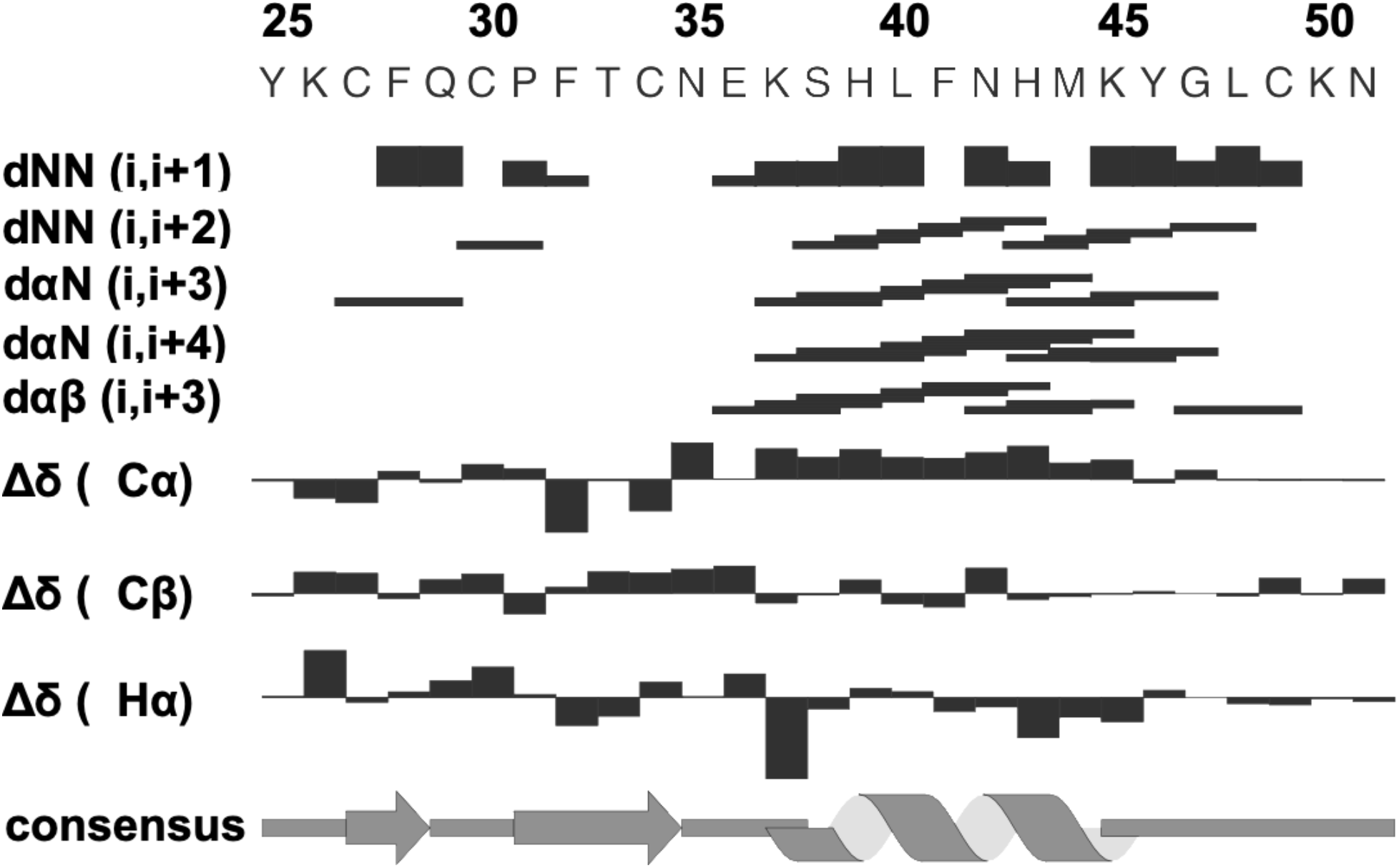
Secondary structure of Z*. The consensus secondary structure is based on NMR short-range NOE distance contacts and chemical shift differences from random coil values, calculated with the program DANGLE (46). The thickness of the horizontal bars denotes the intensity of backbone NOEs.

### The NMR structure of Z* demonstrates a CCHC metal binding site

The ββα secondary structure of Z* (Fig. 4) would seem to preclude a CCHH-type ZNF structure, since this would place the putative ligands C30 in the β-hairpin turn and H39 at the beginning of the α-helix, on opposite ends of the molecule. The preeminent structure prediction algorithm AlphaFold (37,49,50) sheds little light on the matter, as it predicts irregular structure for the Z* domain with a low confidence pLDDT score (Fig. 5A). Sequence conservation within Z* is high (9), and did not allow us to distinguish between the three alternative sets of chelating residues. To determine the correct metal ligands and fold for Z*, we calculated its solution NMR structure using the experimental data in Table 1.

**Table 1.**
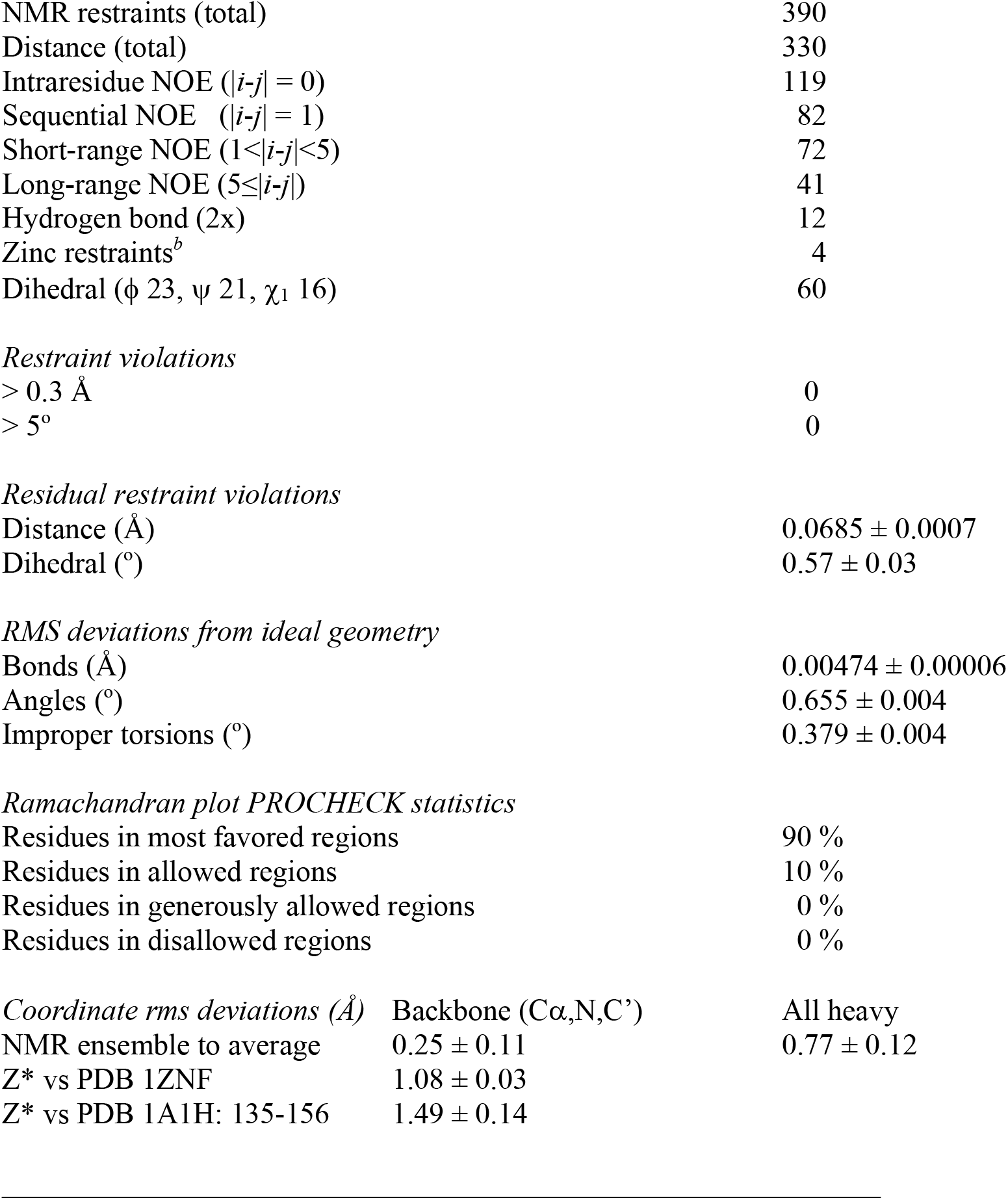
Statistics for the 20 best NMR structures of the Z* domain.

**Figure 5.**
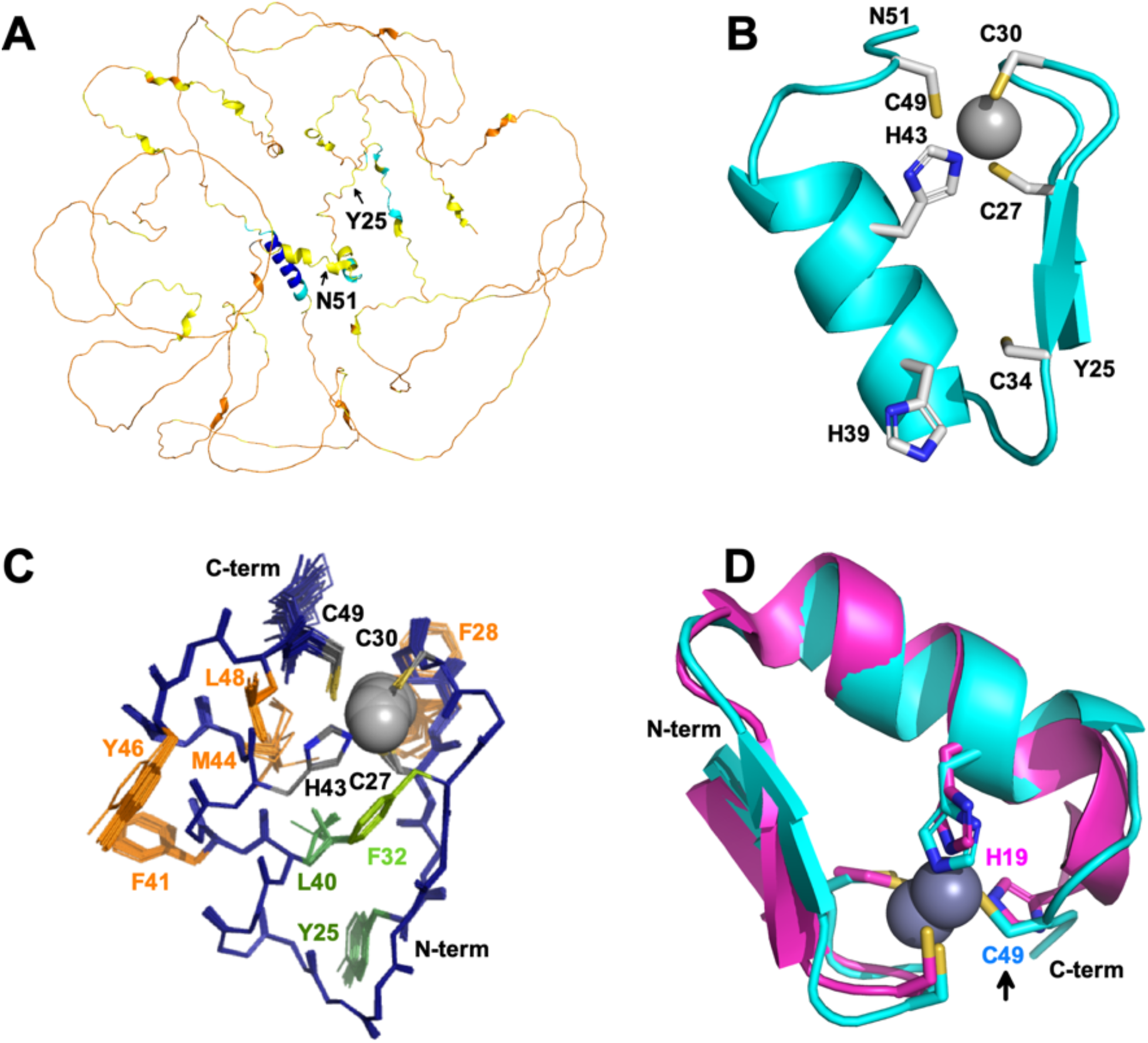
Structure of the Z* domain. **(A)** AlphaFold model for the human ZNF750 protein (PDB CSM accession code AFQ32MQ0F1). The position of the Z* domain from Y25 to N51 is indicated. The model is color-coded according to prediction confidence levels: pLDDT> 90 blue, 90 > pLDDT > 70 cyan, 70 > pLDDT > 50 yellow, 50 < pLDDT orange. Of the 723 a.a. protein, twelve in the α-helix running from 630-642 are predicted with very high confidence. (**B**) Folding topology and Zn^2+^-binding site of the Z* domain based on experimental NMR data (Table 1). The sidechains of Zn^2+^-chelating residues are shown, as well as C34 and H49 which do not participate in metal binding. (**C**) Ensemble of the top NMR structures of Z*. The backbone is shown in dark blue, sidechains of Zn^2+^ ligands in gray, conserved aromatic and hydrophobic residues in green, and additional nonpolar residues that are not part of the conserved consensus in orange. (**D**) Superposition of the NMR structure of Z* from this work with the first structure of a ZNF (73), the classical CCHH domain from *Xfin-31* (PDB code 1ZNF). Note the similarity in overall structures, and in the positioning of the last ligand (H19 in *Xfin-31*, C49 in Z*) relative to the metal.

In the NMR structure calculations, we took great care not to bias the structures by introducing incorrect restraints between the potential ligating residues and Zn^2+^. We calculated an initial set of NMR structures using only dihedral and intra-protein distance restraints. The initial structures showed residues C27, C30, H43 and C49 were in proximity and poised to ligate Zn^2+^. By contrast residues C34 and H39 were too far to participate in metal binding. Once correct ligands were identified from the initial structures, distance restraints to the Zn^2+^ atom were included in the final set of NMR structure calculations (Fig. 5). As an additional check of correct ligand assignments, we used 1D NMR to demonstrate that the variant Z*(C34A, H39A), missing the cysteine and histidine not involved in ligation, retains the ability to fold with Zn^2+^ (Fig. 1D).

A ribbon diagram for the NMR structure of Z* closest to the ensemble average is shown in Fig. 5B. The set of 20 lowest energy structures with no violations is displayed in Fig. 5C. The Z* structure is superposed with the *Xfin-31* classical CCHH ZNF in Fig. 5D. The two structures are remarkably close with a backbone RMSD of 1.1 Å. Comparison to other ZNF structures (Fig. 6A), gave similar results with a range of RMSDs between 0.8 and 1.5 Å.

**Figure 6.**
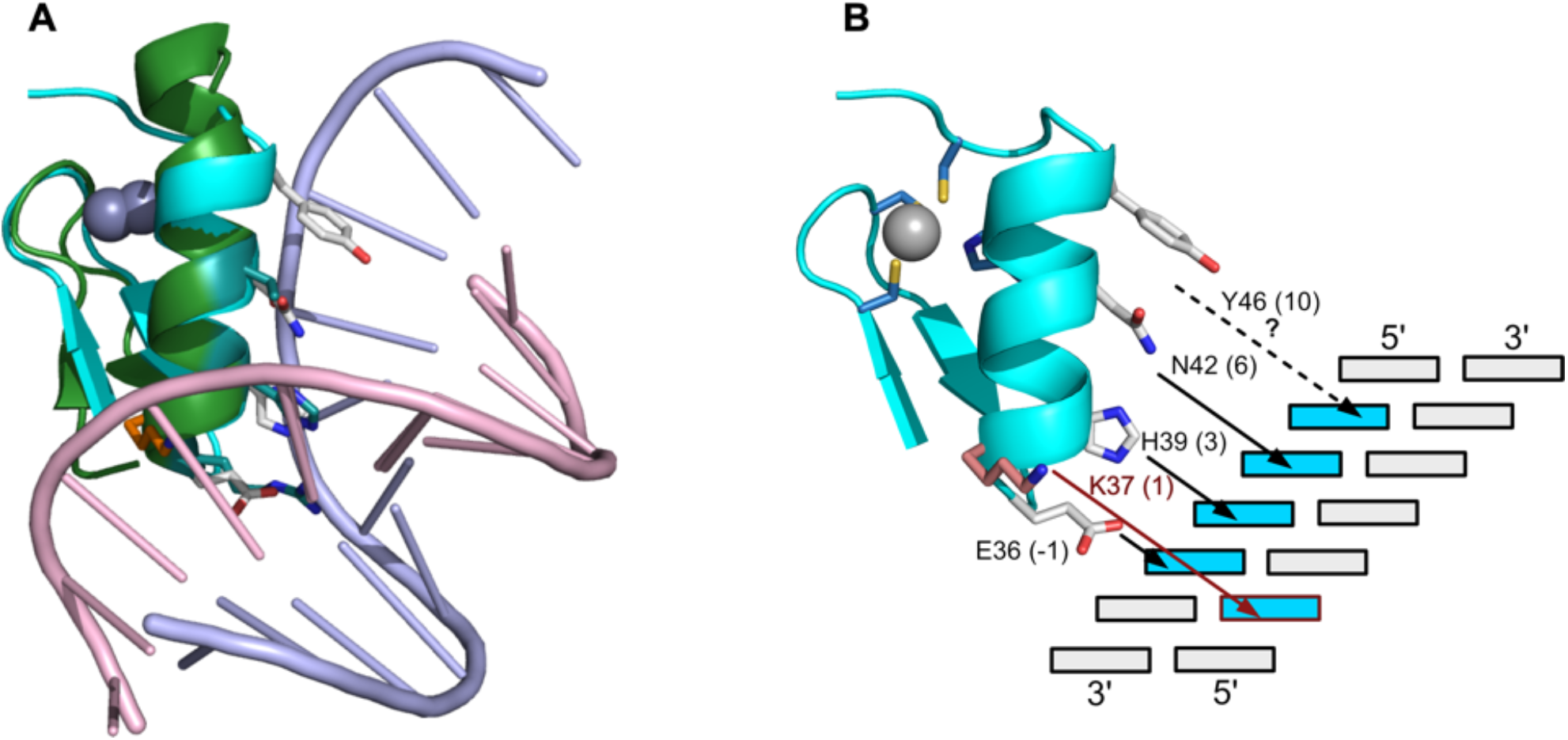
Predicted DNA binding site for Z*. **(A)** Superposition of Z* (cyan ribbon, white sidechains) onto the middle ZNF of *Egr-1* (green) in complex with its target DNA (PDB 4X9J). The 1.4 Å X-ray structure of *Egr-1* (52) has two flanking ZNFs in the DNA complex that are omitted for clarity. The superposition illustrates the similarity of the ZNF structures (RMSD 1.2 Å) and the DNA-binding sidechains of Z* (white) relative to the Erg-1 ZNF. (**B**) Schematic of the predicted DNA-binding residues of Z* based on the ZNF consensus. In addition to the consensus residues, Y46 may bind to the coding strand, and K37 to the noncoding strand.

Compared to *Xfin-31* that is a CCHH-type ZNF, the final half-turn of α-helix in Z* is disrupted to allow C49 to become the last ligand in a tetrahedral CCHC coordination site for Zn^2+^ (Fig. 1D). The first ligand C27 comes from stand β¹, the second C30 from the reverse turn between the two β-strands, and the third H43 from the middle of the α-helix (Fig. 5A) – all features of the classical CCHH ZNF structure (Fig. 1C-E). To a large extent the backbone structure of the CCHC domain Z*, is very much like that of a CCHH ZNF except that the last ligand is a cysteine rather than a histidine. Structural similarity extends to the residue following the second cysteine ligand, P31, having unusual backbone dihedral angles (39) that account for most of the deviations of Z* from Ramachandran ideality (Table 1). The structural similarity is even more striking for the sidechains around the Zn^2+^ ion (Fig. 5D). The first three conserved ligands give an RMSD of 0.36 Å over 15 equivalent heavy atoms, compared to 1.08 Å for the backbone atoms of the entire domain. For the non-conserved fourth ligand, the atoms Cα, Cβ, Sγ of C49 in Z*, essentially trace the Nε1, Cε1, and Nε2 aromatic ring atoms of H23 in *Xfin-31*, to result in structurally if not chemically nearly equivalent coordination sites (arrow in Fig. 5D).

Of the Cys and His residues not involved in metal coordination (Fig. 5D), H39 is likely involved in DNA binding (see below). The cysteine that does not participate in metal coordination, C34, points to the interior of the structure with its sulfhydryl group possibly forming a hydrogen bond with the backbone carbonyl of the last residue before the start of the α-helix, E36.

The three residues Y25, F32, and L40 (green in Fig. 5C) correspond to the conserved hydrophobic core at the base of the V-like wedge between the β-hairpin and α-helix in classical ZNFs. Although, F32 (light green in Fig. 5C) is strictly not-conserved (Fig. 1B,G), it serves the consensus structural role (Fig. 1B). In addition to the classic hydrophobic core, Z* appears to have a second hydrophobic ridge that stabilizes the structure comprised of residues F28, F41, M44, Y46 and L48 (orange in Fig. 5C).

### The structure of Z* identifies DNA-binding residues and provides a structural context for cancer-associated mutants

In classical CCHH ZNFs, the DNA binding site is formed by residues -1, 3, and 6, where the numbering corresponds to the start of the α-helix (24,51). Given the structure of Z* is so close to that of classical ZNFs, we superposed it onto the best-available 1.4 Å-resolution X-ray structure of a ZNF-DNA complex (52), to gain insights into DNA-binding mode. Note that the X-ray structure of *Egr-1* (PDB code 4X9J) has a tandem of three ZNFs bound to DNA. For clarity only the middle finger is shown in Fig. 6A. The superposition and sequence rules for ZNFs (24,51) allow us to identify the putative DNA-ligating residues in Z* as E36 (−1), H39 (3) and N42 (6) (Fig. 6B). The sidechain of Y46 at the 10 position of Z* also seems poised for interactions with the base above that recognized by the 6 position (Fig. 6B), a pattern found in some ZNFs with a canonical DNA binding mode (51). A common feature of the DNA recognition motif in canonical ZNFs, is that the residue in position 2 recognizes a base on the noncoding strand (Fig. 6B), to create a 4 base pair overlapping subsite in transcription factors with multiple tandem ZNFs (24). In Z*, the 2 position is occupied by S38 the sidechain of which seems too small to reach the opposite strand. Rather, K37 in the 1 position appears ideally positioned to interact with the noncoding strand. The details of the DNA-binding mechanism for Z* will require the structure of a DNA complex, which may prove difficult for ZNF750, since it has only a single ZNF domain.

Genome-wide association studies (GWAS) of cancer patients have identified several somatic missense mutations in the ZNF750 transcription factor, with some of these specifically in the Z* domain (Table 2). The mutations were compiled from the literature (14) and from the CBioPortal (53) database (http://www.cbioportal.org). We caution that the significance of these somatic mutations is unknown, since a higher mutation rate due to impaired DNA repair is a hallmark of cancers in general (54,55). Of the currently catalogued cancer-associated mutations in Z*, all involve structural residues rather than amino acids that could participate in the DNA binding site (Table 2). This includes the C49R mutation that replaces the cysteine we established in this work is the fourth ligand for Zn^2+^. In addition to missense mutations, there are frameshift and nonsense mutations that affect the function of the transcription factor. In the familial SLDP disease that led to the discovery of the ZNF750 gene (9), an autosomal dominant frameshift mutation deletes everything past residue 19 of the 723-residue protein, including the Z* domain.

**Table 2.** Cancer-associated somatic missense mutations in the Z* domain of ZNF750.

| **Mutation** | **ID<sup>a</sup> or reference** | **Cancer type** | **Site role in structure** |
| --- | --- | --- | --- |
| Y25D | TCGA-Z6-A8JE | Esophageal squamous cell | conserved aromatic |
| K26E | TCGA-D3-A3C7 | Cutaneous melanoma | α-helix C-cap |
| F28L | TCGA-TN-A7HL | Head and neck squamous | β-hairpin turn aromatic |
| F28Y | TCGA-TN-A7HL | Head and neck squamous | β-hairpin turn aromatic |
| P31S | TCGA-E7-A677 | Bladder urothelial | β-hairpin turn proline |
| F32L | (14) | Esophageal squamous cell | conserved aromatic |
| L40I | TCGA-BS-A0UV | Uterine endometrioid | conserved hydrophobic |
| L40V | TCGA-DK-A3WW | Bladder urothelial | conserved hydrophobic |
| M44R | (14) | Esophageal squamous cell | hydrophobic in core |
| C49R | TCGA-BB-7870 | Head and neck squamous | Loss of 4<sup>th</sup> Zn<sup>2+</sup>-ligand |

<sup>a</sup>Sample IDs are from for the CBioPortal (53) (http://www.cbioportal.org) for the ZNF750 entry.

## DISCUSSION

As the number of protein structures has grown, the trend in molecular biology has been to address experimentally uncharacterized proteins by modeling. Certainly, modeling is much less costly and difficult than experiments. If protein structures could be accurately predicted, the basic science of structural biology would be made redundant to increasingly translational biological research. The present results on Z* indicate abandonment of experimental structural biology (37,49,50) is not yet warranted. The Z* domain is classified as degenerate by the UniProt database because its sequence does not conform to the classical CCHH ZNF consensus (34,35). Besides Z*, we have recently determined that the sixth ZNF domain of ZNF711, also classified as degenerate (UniProt accession code Q9Y462), folds in the presence of Zn^2+^ (A.J.R. and A.T.A., in preparation). We are aware of another two ZNF domains classified as degenerate by UniProt (accession codes Q9P243 domain 14, P17041 domain 7) that have NMR structures in the Protein Databank (PDB codes 2RV1 and 2EPC).

The AlphaFold method that performed the best at the last few biennial CASP tournaments (49,56), predicts at low confidence a structure for Z* that differs from our experimental NMR structure (Fig. 5A). About 42% of residues in human proteins are predicted by AlphaFold with low confidence, defined by a pLDDT score lower than 70 (57). The inference for these regions is that they are dynamic, either intrinsically disordered or becoming structured as part of a complex (57). If true, AlphaFold would not only predict protein structure but flexibility. The structure of Z* is extremely stable with a melting point above 80 °C. S^2^ order parameters calculated from NMR chemical shifts indicate the structure is rigid except for the last two residues in the sequence, in agreement with the tight clustering of conformers in the NMR ensemble. We suspect that the low confidence of the AlphaFold prediction for Z* has less to do with dynamics than with the unfamiliarity of its sequence, that make the domain underrepresented in sequence and structure databases. ZNFs have more structural variability than typical protein folds (42,43,58). A danger of Big Science genomic conformity is that it can paint protein structures with a large brush. This might miss interesting consensus exceptions that offer the best chance to discover novel insights into the versatility of protein structures by defying predictions based on recycled experimental results.

The first clues that the Z* domain is a functional ZNF came from experiments demonstrating the double-mutants ZNF750(C27A,C30A), ZNF750(H39A, H43A) or the quadruple mutant ZNF750(C27A,C30A, H39A, H43A) abrogated the ability of the transcription factor to induce late terminal epidermal differentiation through its induction of the expression of the KLF4 transcription factor (3,4). Mutagenesis can lead to ambiguous results since a missense mutation can affect structural or functional parts of a protein. Indeed, based on our present work the H39A mutation interferes with DNA-binding rather than Zn^2+^-chelation (Fig. 5B, 6B). Because of the assumption of a CCHH ZNF, and because the putative ligating residues were mutated in pairs, the mutagenesis experiments missed that the Z* domain has a CCHC (C27, C30, H43, C49) rather than the expected CCHH coordination site. The CCHH arrangement (C27, C30, H39, H43) is disfavored by the prohibitively short 8-residue spacing between the second and third ligands (Fig. 1F, 5B).

There are two conceivable ways a ZNF domain could fold despite sequence differences from consensus. The first is a rearrangement of the structure to preserve the character of the metal binding site. The second is to allow differences in the Zn^2+^ ligands, while preserving the folding motif. The second alternative is used by the Z* domain. Similarly, the previously described examples of degenerate ZNFs with structures in the PDB (codes 2RV1 and 2EPC), both have the prototypical ββα fold. This suggests selective pressure to maintain the ββα structural motif determined by the spacing between ligands and conserved positions forming the hydrophobic core. By contrast constraints on the identity of the Zn^2+^ binding ligands appear laxer, as long as the ββα-fold foundation for the coordination site is preserved.

From a structural point of view, the Z* domain is just like a classical CCHH-type ZNF, except that the last Zn^2+^ ligand histidine ligand is replaced by a cysteine (Fig. 5D). The Zn^2+^-ligating residues are on the opposite side of the α-helix that determines the DNA binding-site of CCHH-type ZNFs (Fig. 6B). In Z*, neither Zn^2+^-binding affinity (Fig. 2B) nor stability (Fig. 2A) are perceptibly affected by the switch of the last ligand from a histidine to a cysteine. This suggests that the association of CCHH-type ZNFs with DNA-binding functions is due to gene duplication and divergent evolution. The structure and stability properties of these motifs appear equally well supported whether a histidine or cysteine is the last ligand for Zn^2+^.

There is a CCHC family of ZNF domains that has a sequence spacing C-X_2_-C-X_4_-H-X4-C. This family typically functions as an RNA-binding module in RNA metabolism processes (27,29,35). Member of the family usually have a ‘zinc knuckle’ structure, where the short four-residue segment between the second cysteine and histidine ligands, truncates the ‘finger’ to a knuckle. We think it is useful to distinguish this knuckle-CCHC family from the finger-CCHC family exemplified by the Z* domain, since the latter have structures very similar to the classical ZNFs. Besides Z*, other examples of finger-CCHC domains exist in the proteins NEMO (59), Fog, MOZ, and U-shaped (60). In these examples the CCHC-finger motif is used in protein-protein interactions. By contrast the seventh domain of ZNF32 (PDB 2EPC) is probably a DNA-binding CCHC-finger, like Z*, since it occurs in a transcription factor.

We anticipate there are substantially more CCHC-fingers yet to be discovered. Some of the CCHH ZNFs classified by UniProt as degenerate, occur in transcription factors and have a cystine residue replacing the last histidine (examples include: Q6NUN9-1, P1004-2, Q2M1K9-1, Q9P243-2, Q9NQX0-1, P52739-4 Q86UP3-15, Q86UP3-19; where the UniProt accession code is followed by the ZNF domain designated as degenerate). Other degenerate CCHH ZNFs that appear to have only the first three of the four ligands, can be ‘rescued’ by extending the domain boundary towards the C-terminus to bring in a nearby cysteine. Given that the end of the ZNF α-helix is often frayed or followed by an irregular segment, the cysteine likely corresponds to the fourth ligand (examples include: Q9NQX0-4, Q9NU63-7, Q8TD23-3). The folding status of these domains will need to be confirmed experimentally.

ChIP-Seq data demonstrated that a segment of the ZNF750 transcription factor near the Z* domain binds the progenitor genes it represses and the differentiation genes it activates, through a 5’-CCNNAGGC-3’ DNA consensus sequence (3). The ChIP-Seq method uses chromatin immunoprecipitation together with DNA sequencing, for genome-wide profiling of protein DNA binding-sites (61). The quadruple ZNF750 (C27A, C30A, H39A, H43A) mutant described previously, significantly decreased DNA binding (3), indicating the Z* motif is required for this function. The DNA-binding site prediction server for ZNFs (http://zf.princeton.edu) (62), suggests the Z* domain has a preference for the DNA sequence pattern (G/C)G(C/G), with only the central G of the triplet exhibiting a strong propensity. The prediction is in good agreement with our NMR structure, since H39 in the 3 position of the Z* binding site would have a strong preference for guanine in the central position of the DNA triplet (51). The predicted (G/C)G(C/G) DNA-binding pattern for Z*, agrees with the end of the CCNNA<u>GGC</u> consensus DNA-sequence determined by the ChIP-Seq data (3). However, a single ZNF domain can typically recognize only a single DNA triplet (24,51), not the eight base-pair consensus implied by the ChIP-Seq data. To recognize larger DNA segments, transcription factors typically use multiple ZNFs domains in tandem (24,63). The typical folding independence of the ZNF domains has been used to design ZNF-nuclease gene editors (24,51), as an early alternative to the CRISPR/CAS9 system (64).

In this regard, ZNF750 is an extremely rare and unusual example of a transcription factor with a single ZNF domain. We are aware of only one other such example, the single ZNF domain from the GAGA transcription factor (23). With the GAGA transcription factor, NMR structural studies showed that two basic regions from the N-terminus of the protein adopt α-helical structures that augment the site-specific DNA binding modulated by the single ZNF domain (23). Like the GAGA transcription factor, ZNF750 has basic regions upstream of the Z* domain (light blue in Fig. 1F). In ZNF750 the N-terminal basic region includes the sequence RKPKK (underlined in Fig. 1F) that is a signature nuclear localization signal (NLS). However, there are additional NLS motifs at the C-terminus of ZNF750 (Fig. 1A), and it was shown that the N-terminal NLS was dispensable for nuclear localization (5). The N-terminal basic region could thus be part of an extended DNA-binding site that includes the Z* domain, like in the GAGA transcription factor. Alternatively, immunoprecipitation and mutagenesis studies showed that ZNF750 interacts with the chromatin regulatory proteins KDM1A, RCOR1, CTPB1 and CTPB1, through two PLNLS amino acid sequence motifs in the 504-563 region (Fig. 1A) of the protein (3). Moreover, mutagenesis of the PLNLS motifs affected DNA-binding (3). The chromatin regulatory proteins that interact with ZNF750 are transcriptional regulators with inherent DNA-binding functions. Thus, it is possible ZNF750 does not bid DNA on its own but only as part of heterooligomeric complexes that extend the binding site beyond that provided by Z* alone.

Taken together, the structure of Z* sheds new light on the role of this domain in the function and pathology of the ZNF750 transcription factor. There is clearly more data needed on the structural properties of the full-length protein, its interactions with DNA, and with cognate protein binding partners. Given the many unusual features of this transcription factor we expect these aspects will defy predictions and will need experimental data to establish mechanisms unequivocally.

## EXPERIMENTAL PROCEDURES

### Materials

The 27-residue peptides Z* and Z*(C34A, H39A), corresponding to the fragment 25-51 of human ZNF750 (UniProt Q32MQ0), were synthesized by AAPPTec (Louisville, KY). Both peptides had their termini blocked by N-acetylation and C-amidation, were ≥ 90% pure by HPLC, and had molecular weights by mass spectrometry within 3 Daltons of those theoretically expected. ZnSO_4_•H_2_O (purity of ≥ 99.9%) was from Sigma (St. Louis, MO).

### Optical spectroscopy and Zn^2+^ affinity

Z* concentrations were determined via a BCA assay (65) according to the manufacturer’s instructions (Thermo Fisher Scientific; Waltham, MA) on an Ultrospec 8000 UV-Vis spectrophotometer (Thermo Fisher).

Concentrations from the BCA assay were used to determine molar extinction coefficients at 280 nm of 2,954 M^-1^ cm^-1^ for unfolded Z* in the absence of Zn^2+^, and 2,978 M^-1^ cm^-1^ for folded Z* in the presence of a 1:1.25 ratio of Zn^2+^.

CD data were collected using an Applied Photophysics Chirascan V100 Spectrometer (Surrey, UK) using a 1 mm cuvette. The Z* concentration was 124 μM in 10 mM NaPO_4_ buffer, pH 7. CD spectra were collected between 190-250 nm, using a 1 nm bandwidth, a 1 nm step size, and 5 s/point data averaging for an approximate total scan time of 5 minutes. Thermal melt experiments were collected from 20 ºC to 96 ºC in 4-degree increments, with the same parameters except using 0.75 s/point data averaging, for a total scan time of 72 seconds.

For Zn^2+^-binding experiments 22 mM EGTA together with ZnSO_4_ concentrations of 0.1-200 μM were added to 133 µM Z*. The fraction of Zn^2+^-bound protein, Φ, was calculated from

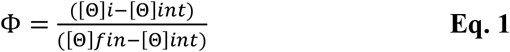

where [Θ]*i* is the measured ellipticity at 200 nm for a given Zn^2+^ concentration, and [Θ]*int* and [Θ]*fin* are the plateau values for free and fully-bound Z*, respectively (66). Free Zn^2+^ concentrations were calculated with the WEBMACX program (40,67). The binding data were modeled according to a sigmoidal function:

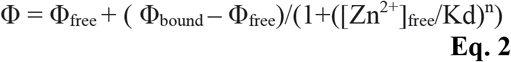

where the fitted parameters were the dissociation constant K_d_, the Hill coefficient n, and the plateau fraction bound values (Φ) at low and high [Zn^2+^]_free_ concentrations (Fig. 2B).

### NMR spectroscopy

To establish folding in the presence of (Fig. 2C,D), 1D ^1^H-NMR experiments with.5 mM samples of Z* and Z*(C34A, H39A) were done at 27 °C on a Bruker Avance 500 MHz spectrometer. PFG-diffusion experiments established that the Zn^2+^-bound domain had an R_h_ of 8.1 ± 1.2 Å, consistent with a monomer (68).

All 2D NMR experiments used for assignments and structure calculations were done at 15 °C on a Bruker Avance 600 MHz instrument equipped with a TCI cryogenic probe, that is part of the Francis Bitter Magnet Lab of MIT. A sample for 2D NMR experiments in H_2_O had 1.5 mM Z* and 2.4 mM ZnSO_4_, in 10 mM NaPO_4_ buffer. For experiments in D_2_O the sample had 3.0 mM Z*, 4.8 mM ZnSO_4_, and 20 mM NaPO_4_ buffer. Both samples had a measured pH of 5.9.

The experiments recorded on the H_2_O sample included 70 ms mixing time TOCSY (*mlevgpphw5*), 200 ms mixing time NOESY (*noesygpphw5*), and a natural abundance ^15^N-sofast-HMQC (*sfhmqcf3gpph*). The names in parentheses are the Bruker pulse programs for the experiments. Experiments on the D_2_O sample included 50 and 200 ms mixing time NOESY, DQF-COSY (*cosydfph*), ECOSY (*ecos3nph*), and natural abundance ^13^C-HSQC (*hsqcetgpsi2*). Most of the assignments were obtained from a sequential walk using the NOESY and TOCSY spectra recorded in H_2_O (Fig. 3A). The natural abundance ^1^H-^13^C HSQC spectrum (Fig. 3B) proved particularly useful to confirm ^1^H aliphatic assignments and to extend these to the bonded carbon resonances. The ^15^N-sofast-HMQC experiment (Fig. 3C) was used to extend amide proton assignments to their bonded nitrogens. Experiments in D_2_O were used for aromatic assignments, and to observe Hα resonances without interference from solvent. The extent of assigned resonances was ^1^H (93%), ^13^C (83%, excluding C’), and ^15^N (77%). Stereospecific assignments were obtained from ECOSY and short mixing time (50 ms) NOESY experiments (69).

### NMR Structure calculations

The restraints used for NMR structure calculations are summarized in Table I. Distance restraints were obtained from NOESY crosspeak intensities calculated with the program CcpNmr Analysis 2.5 (70). Torsional restraints were calculated from ^1^H_N_, ^1^Hα ^13^Cα, ^13^Cβ, and ^15^N chemical shifts using the program TALOS-N (71). An initial set of 20 lowest energy structures without violations was calculated from a starting set of 100 random conformation with the program XPLOR-NIH (72) to identify the Zn^2+^ ligands. These initial structures showed an anti-parallel ββα structure where C27, C30, H43, and C49 were poised to bind Zn^2+^ but C34 and H39 were too far to participate in the coordination site. Based on these initial structures, distance constraints were included between Zn^2+^ and the aforementionedligands, and hydrogen bond (H-bond) restraints were included for amide protons protected from solvent exchange.

Two distance restraints were used per H-bond (1.5-2.5 Å for NH-O and 2.5-3.5 Å for N-O) to enforce linearity. An additional seven distance restraints were included between the protein and the Zn^2+^ atom: 2.33-2.37 Å for Zn^2+^-Sγ and *r*_Zn-Cβ_ = 3.25-3.51 Å for Zn^2+^-Sγ‐ for each of the three cysteines. An ambiguous restraint of 1.0-3.1 Å was set between Zn^2+^ and either the Nε1 or Nε2 atom of H43. In all the final structures the Zn^2+^ was bonded to the Nε2 atom of H43. With all metal restraints removed, the structures retained the ββα fold, and gave an all-atom RMSD to the final structures of 0.37 ± 0.05 Å. Thus, the Z* structure is not constrained by the Zn^2+^ contacts alone but by the totality of dihedral and distance restraints.

### Database accession numbers

NMR assignments and chemical shift values for the Z* domain were deposited in the BMRB under accession code 51951. Coordinates for the 20 lowest energy structures with no violations, together with the NOE distance and dihedral restraints used for NMR structure calculations, were deposited in the PDB under accession code 8SXM.

## ACKNOWLEDGEMENTS

We thank Prof. Charles Giardina for useful discussion, Prof. Carolyn Teschke for use of her CD spectrophotometer, Prof. Nathan Alder for use of his UV-Vis spectrophotometer, and Prof Mei Hong for use of her group’s solution 600 MHz instrument during the sabbatical of A.T.A. The NMR experiments used equipment at the MIT-Harvard Center for Magnetic Resonance, which is supported by the P41 grant GM132079. M.H. is partially supported by NIH grant AG059661.

## REFERENCES

1. Lambert, S. A., Jolma, A., Campitelli, L. F., Das, P. K., Yin, Y., Albu, M., Chen, X., Taipale, J., Hughes, T. R., and Weirauch, M. T. (2018) The Human Transcription Factors. Cell 172, 650–665

2. Emerson, R. O., and Thomas, J. H. (2009) Adaptive evolution in zinc finger transcription factors. PLoS Genet 5, e1000325

3. Boxer, L. D., Barajas, B., Tao, S., Zhang, J., and Khavari, P. A. (2014) ZNF750 interacts with KLF4 and RCOR1, KDM1A, and CTBP1/2 chromatin regulators to repress epidermal progenitor genes and induce differentiation genes. Genes Dev 28, 2013–2026

4. Sen, G. L., Boxer, L. D., Webster, D. E., Bussat, R. T., Qu, K., Zarnegar, B. J., Johnston, D., Siprashvili, Z., and Khavari, P. A. (2012) ZNF750 is a p63 target gene that induces KLF4 to drive terminal epidermal differentiation. Dev Cell 22, 669–677

5. Cohen, I., Birnbaum, R. Y., Leibson, K., Taube, R., Sivan, S., and Birk, O. S. (2012) ZNF750 is expressed in differentiated keratinocytes and regulates epidermal late differentiation genes. PLoS One 7, e42628

6. Zarnegar, B. J., Webster, D. E., Lopez-Pajares, V., Vander Stoep Hunt, B., Qu, K., Yan, K. J., Berk, D. R., Sen, G. L., and Khavari, P. A. (2012) Genomic profiling of a human organotypic model of AEC syndrome reveals ZNF750 as an essential downstream target of mutant TP63. Am J Hum Genet 91, 435–443

7. Lopez-Pajares, V., Yan, K., Zarnegar, B. J., Jameson, K. L., and Khavari, P. A. (2013) Genetic pathways in disorders of epidermal differentiation. Trends Genet 29, 31–40

8. Wikramanayake, T. C., Stojadinovic, O., and Tomic-Canic, M. (2014) Epidermal Differentiation in Barrier Maintenance and Wound Healing. Adv Wound Care (New Rochelle) 3, 272–280

9. Birnbaum, R. Y., Zvulunov, A., Hallel-Halevy, D., Cagnano, E., Finer, G., Ofir, R., Geiger, D., Silberstein, E., Feferman, Y., and Birk, O. S. (2006) Seborrhea-like dermatitis with psoriasiform elements caused by a mutation in ZNF750, encoding a putative C2H2 zinc finger protein. Nat Genet 38, 749–751

10. Birnbaum, R. Y., Hayashi, G., Cohen, I., Poon, A., Chen, H., Lam, E. T., Kwok, P. Y., Birk, O. S., and Liao, W. (2011) Association analysis identifies ZNF750 regulatory variants in psoriasis. BMC Med Genet 12, 167

11. Yang, C. F., Hwu, W. L., Yang, L. C., Chung, W. H., Chien, Y. H., Hung, C. F., Chen, H. C., Tsai, P. J., Fann, C. S., Liao, F., and Chen, Y. T. (2008) A promoter sequence variant of ZNF750 is linked with familial psoriasis. J Invest Dermatol 128, 1662–1668

12. Shao, S., Tsoi, L. C., Swindell, W. R., Chen, J., Uppala, R., Billi, A. C., Xing, X., Zeng, C., Sarkar, M. K., Wasikowski, R., Jiang, Y., Kirma, J., Sun, J., Plazyo, O., Wang, G., Harms, P. W., Voorhees, J. J., Ward, N. L., Ma, F., Pellegrini, M., Merleev, A., Perez White, B. E., Modlin, R. L., Andersen, B., Maverakis, E., Weidinger, S., Kahlenberg, J. M., and Gudjonsson, J. E. (2021) IRAK2 Has a Critical Role in Promoting Feed-Forward Amplification of Epidermal Inflammatory Responses. J Invest Dermatol 141, 2436–2448

13. Xu, C., Yang, H. L., Yang, Y. K., Pan, L., and Chen, H. Y. (2022) Zinc-finger protein 750 mitigates malignant biological behavior of oral CSC-like cells enriched from parental CAL-27 cells. Oncol Lett 23, 28

14. Takahashi, M., Hosomichi, K., Nakaoka, H., Sakata, H., Uesato, N., Murakami, K., Kano, M., Toyozumi, T., Matsumoto, Y., Isozaki, T., Sekino, N., Otsuka, R., Inoue, I., and Matsubara, H. (2022) Biased expression of mutant alleles in cancer-related genes in esophageal squamous cell carcinoma. Esophagus 19, 294–302

15. Du, Y., Lv, G., Jing, C., Liu, J., and Liu, J. (2020) ZNF750 inhibits the proliferation and invasion of melanoma cells through modulating the Wnt/b-catenin signaling pathway. Folia Histochem Cytobiol 58, 255–263

16. North, J. P. (2021) Molecular Genetics of Sebaceous Neoplasia. Surg Pathol Clin 14, 273–284

17. Montanaro, M., Agostini, M., Anemona, L., Bonanno, E., Servadei, F., Finazzi Agro, E., Asimakopoulos, A. D., Ganini, C., Cipriani, C., Signoretti, M., Bove, P., Rugolo, F., Imperiali, B., Melino, G., Mauriello, A., and Scimeca, M. (2023) ZNF750: A Novel Prognostic Biomarker in Metastatic Prostate Cancer. Int J Mol Sci 24

18. Butera, A., Cassandri, M., Rugolo, F., Agostini, M., and Melino, G. (2020) The ZNF750-RAC1 axis as potential prognostic factor for breast cancer. Cell Death Discov 6, 135

19. An, G., Feng, L., Hou, L., Li, X., Bai, J., He, L., Gu, S., and Zhao, X. (2022) A bioinformatics analysis of zinc finger protein family reveals potential oncogenic biomarkers in breast cancer. Gene 828, 146471

20. Cassandri, M., Butera, A., Amelio, I., Lena, A. M., Montanaro, M., Mauriello, A., Anemona, L., Candi, E., Knight, R. A., Agostini, M., and Melino, G. (2020) ZNF750 represses breast cancer invasion via epigenetic control of prometastatic genes. Oncogene 39, 4331–4343

21. Cassandri, M., Smirnov, A., Novelli, F., Pitolli, C., Agostini, M., Malewicz, M., Melino, G., and Raschella, G. (2017) Zinc-finger proteins in health and disease. Cell Death Discov 3, 17071

22. Lee, M. S., Gottesfeld, J. M., and Wright, P. E. (1991) Zinc is required for folding and binding of a single zinc finger to DNA. FEBS Lett 279, 289–294

23. Omichinski, J. G., Pedone, P. V., Felsenfeld, G., Gronenborn, A. M., and Clore, G. M. (1997) The solution structure of a specific GAGA factor-DNA complex reveals a modular binding mode. Nat Struct Biol 4, 122–132

24. Klug, A. (2010) The discovery of zinc fingers and their applications in gene regulation and genome manipulation. Annu Rev Biochem 79, 213–231

25. Miller, J., McLachlan, A. D., and Klug, A. (1985) Repetitive zinc-binding domains in the protein transcription factor IIIA from Xenopus oocytes. EMBO J 4, 1609–1614

26. Berg, J. M., and Shi, Y. (1996) The galvanization of biology: a growing appreciation for the roles of zinc. Science 271, 1081–1085

27. Michalek, J. L., Besold, A. N., and Michel, S. L. (2011) Cysteine and histidine shuffling: mixing and matching cysteine and histidine residues in zinc finger proteins to afford different folds and function. Dalton Trans 40, 12619–12632

28. Fu, M., and Blackshear, P. J. (2017) RNA-binding proteins in immune regulation: a focus on CCCH zinc finger proteins. Nat Rev Immunol 17, 130–143

29. Wang, Y., Yu, Y., Pang, Y., Yu, H., Zhang, W., Zhao, X., and Yu, J. (2021) The distinct roles of zinc finger CCHC-type (ZCCHC) superfamily proteins in the regulation of RNA metabolism. RNA Biol 18, 2107–2126

30. Ravasi, T., Huber, T., Zavolan, M., Forrest, A., Gaasterland, T., Grimmond, S., Hume, D. A., Group, R. G., and Members, G. S. L. (2003) Systematic characterization of the zinc-finger-containing proteins in the mouse transcriptome. Genome Res 13, 1430–1442

31. Branden, C., and Tooze, J. (1998) Introduction to Protein Structure, Garland Science, New York

32. Lachenmann, M. J., Ladbury, J. E., Qian, X., Huang, K., Singh, R., and Weiss, M. A. (2004) Solvation and the hidden thermodynamics of a zinc finger probed by nonstandard repair of a protein crevice. Protein Sci 13, 3115–3126

33. Parraga, G., Horvath, S., Hood, L., Young, E. T., and Klevit, R. E. (1990) Spectroscopic studies of wild-type and mutant “zinc finger” peptides: determinants of domain folding and structure. Proc Natl Acad Sci U S A 87, 137–141

34. The_UniProt_Consortium. (2023) UniProt: the Universal Protein Knoledgebase in 2023. Nucleic Acids Res 51, D523–D531

35. Aceituno-Valenzuela, U., Micol-Ponce, R., and Ponce, M. R. (2020) Genome-wide analysis of CCHC-type zinc finger (ZCCHC) proteins in yeast, Arabidopsis, and humans. Cell Mol Life Sci 77, 3991–4014

36. Jumper, J., Evans, R., Pritzel, A., Green, T., Figurnov, M., Ronneberger, O., Tunyasuvunakool, K., Bates, R., Zidek, A., Potapenko, A., Bridgland, A., Meyer, C., Kohl, S. A. A., Ballard, A. J., Cowie, A., Romera-Paredes, B., Nikolov, S., Jain, R., Adler, J., Back, T., Petersen, S., Reiman, D., Clancy, E., Zielinski, M., Steinegger, M., Pacholska, M., Berghammer, T., Bodenstein, S., Silver, D., Vinyals, O., Senior, A. W., Kavukcuoglu, K., Kohli, P., and Hassabis, D. (2021) Highly accurate protein structure prediction with AlphaFold. Nature 596, 583–589

37. Lewis, T. (2023) The AI Biologist: DeepMind’s Demis Hassabis explains how artificial intelligence solved one of the biggest problems in biology. Scientific American 328, 28–30

38. Tremblay, C., Bedard, M., Bonin, M. A., and Lavigne, P. (2016) Solution structure of the 13th C2H2 Zinc Finger of Miz-1. Biochem Biophys Res Commun 473, 471–475

39. Simpson, R. J., Cram, E. D., Czolij, R., Matthews, J. M., Crossley, M., and Mackay, J. P. (2003) CCHX zinc finger derivatives retain the ability to bind Zn(II) and mediate protein-DNA interactions. J Biol Chem 278, 28011–28018

40. Ivanova, E., Ball, M., and Lu, H. (2008) Zinc binding of Tim10: Evidence for existence of an unstructured binding intermediate for a zinc finger protein. 71, 467–475

41. Kluska, K., Adamczyk, J., and Krezel, A. (2018) Metal binding properties, stability and reactivity of zinc fingers. Coordination Chemistry Reviews 367, 18–64

42. Neuhaus, D. (2022) Zinc finger structure determination by NMR: Why zinc fingers can be a handful. Prog Nucl Magn Reson Spectrosc 130-131, 62–105

43. Padjasek, M., Kocyla, A., Kluska, K., Kerber, O., Tran, J. B., and Krezel, A. (2020) Structural zinc binding sites shaped for greater works: Structure-function relations in classical zinc finger, hook and clasp domains. J Inorg Biochem 204, 110955

44. Matousek, W. M., and Alexandrescu, A. T. (2004) NMR structure of the C-terminal domain of SecA in the free state. Biochim Biophys Acta 1702, 163–171

45. Ramboarina, S., Morellet, N., Fournie-Zaluski, M. C., and Roques, B. P. (1999) Structural investigation on the requirement of CCHH zinc finger type in nucleocapsid protein of human immunodeficiency virus 1. Biochemistry 38, 9600–9607

46. Cheung, M. S., Maguire, M. L., Stevens, T. J., and Broadhurst, R. W. (2010) DANGLE: A Bayesian inferential method for predicting protein backbone dihedral angles and secondary structure. J Magn Reson 202, 223–233

47. Shen, Y., and Bax, A. (2013) Protein backbone and sidechain torsion angles predicted from NMR chemical shifts using artificial neural networks. J Biomol NMR 56, 227–241

48. Berg, J. M. (1988) Proposed structure for the zinc-binding domains from transcription factor IIIA and related proteins. Proc Natl Acad Sci U S A 85, 99–102

49. Callaway, E. (2022) ‘The entire protein universe’: AI predicts shape of nearly every known protein. Nature 608, 15–16

50. Kennedy, D. (2001) Breakthrough of the year. Science 294, 2429

51. Wolfe, S. A., Nekludova, L., and Pabo, C. O. (2000) DNA recognition by Cys2His2 zinc finger proteins. Annu Rev Biophys Biomol Struct 29, 183–212

52. Zandarashvili, L., White, M. A., Esadze, A., and Iwahara, J. (2015) Structural impact of complete CpG methylation within target DNA on specific complex formation of the inducible transcription factor Egr-1. FEBS Lett 589, 1748–1753

53. Gao, J., Aksoy, B. A., Dogrusoz, U., Dresdner, G., Gross, B., Sumer, S. O., Sun, Y., Jacobsen, A., Sinha, R., Larsson, E., Cerami, E., Sander, C., and Schultz, N. (2013) Integrative analysis of complex cancer genomics and clinical profiles using the cBioPortal. Sci Signal 6, pl1

54. Jackson, A. L., and Loeb, L. A. (1998) The mutation rate and cancer. Genetics 148, 1483–1490

55. Bailey, C. G., Gupta, S., Metierre, C., Amarasekera, P. M. S., O’Young, P., Kyaw, W., Laletin, T., Francis, H., Semaan, C., Sharifi Tabar, M., Singh, K. P., Mullighan, C. G., Wolkenhauer, O., Schmitz, U., and Rasko, J. E. J. (2021) Structure-function relationships explain CTCF zinc finger mutation phenotypes in cancer. Cell Mol Life Sci 78, 7519–7536

56. Sippl, M. J. (1999) Who solved the protein folding problem? Structure 7, R81–83

57. Tunyasuvunakool, K., Adler, J., Wu, Z., Green, T., Zielinski, M., Zidek, A., Bridgland, A., Cowie, A., Meyer, C., Laydon, A., Velankar, S., Kleywegt, G. J., Bateman, A., Evans, R., Pritzel, A., Figurnov, M., Ronneberger, O., Bates, R., Kohl, S. A. A., Potapenko, A., Ballard, A. J., Romera-Paredes, B., Nikolov, S., Jain, R., Clancy, E., Reiman, D., Petersen, S., Senior, A. W., Kavukcuoglu, K., Birney, E., Kohli, P., Jumper, J., and Hassabis, D. (2021) Highly accurate protein structure prediction for the human proteome. Nature 596, 590–596

58. Krishna, S. S., Majumdar, I., and Grishin, N. V. (2003) Structural classification of zinc fingers: survey and summary. Nucleic Acids Res 31, 532–550

59. Cordier, F., Vinolo, E., Veron, M., Delepierre, M., and Agou, F. (2008) Solution structure of NEMO zinc finger and impact of an anhidrotic ectodermal dysplasia with immunodeficiency-related point mutation. J Mol Biol 377, 1419–1432

60. Matthews, J. M., Kowalski, K., Liew, C. K., Sharpe, B. K., Fox, A. H., Crossley, M., and MacKay, J. P. (2000) A class of zinc fingers involved in protein-protein interactions biophysical characterization of CCHC fingers from fog and U-shaped. Eur J Biochem 267, 1030–1038

61. Park, P. J. (2009) ChIP-seq: advantages and challenges of a maturing technology. Nat Rev Genet 10, 669–680

62. Persikov, A. V., and Singh, M. (2014) De novo prediction of DNA-binding specificities for Cys2His2 zinc finger proteins. Nucleic Acids Res 42, 97–108

63. Patel, A., Yang, P., Tinkham, M., Pradhan, M., Sun, M. A., Wang, Y., Hoang, D., Wolf, G., Horton, J. R., Zhang, X., Macfarlan, T., and Cheng, X. (2018) DNA Conformation Induces Adaptable Binding by Tandem Zinc Finger Proteins. Cell 173, 221–233 e212

64. Zhang, H. X., Zhang, Y., and Yin, H. (2019) Genome Editing with mRNA Encoding ZFN, TALEN, and Cas9. Mol Ther 27, 735–746

65. Walker, J. M. (2002) The Bicinchoninic Acid (BCA) Assay for Protein Quantitation. in The Protein Protocols Handbook (Walker, J. M. ed.), 2nd Ed., Humana Press, Totowa, NJ. pp 11–15

66. Imanishi, M., Nakaya, T., Morisaki, T., Noshiro, D., Futaki, S., and Sugiura, Y. (2010) Metal-Stimulated Regulation of Transcription by an Artificial Zinc-Finger Protein. 11, 1653–1655

67. MaxChelator.

68. Whitehead, R. D., 3rd, Teschke, C. M., and Alexandrescu, A. T. (2022) Pulse-field gradient nuclear magnetic resonance of protein translational diffusion from native to non-native states. Protein Sci 31, e4321

69. Case, D. A., Dyson, H. J., and Wright, P. E. (1994) Use of chemical shifts and coupling constants in nuclear magnetic resonance structural studies on peptides and proteins. Methods Enzymol 239, 392–416

70. Vranken, W. F., Boucher, W., Stevens, T. J., Fogh, R. H., Pajon, A., Llinas, M., Ulrich, E. L., Markley, J. L., Ionides, J., and Laue, E. D. (2005) The CCPN data model for NMR spectroscopy: development of a software pipeline. Proteins 59, 687–696

71. Shen, Y., and Bax, A. (2015) Protein structural information derived from NMR chemical shift with the neural network program TALOS-N. Methods Mol Biol 1260, 17–32

72. Schwieters, C. D., Kuszewski, J. J., Tjandra, N., and Clore, G. M. (2003) The Xplor-NIH NMR molecular structure determination package. J Magn Reson 160, 65–73

73. Lee, M. S., Gippert, G. P., Soman, K. V., Case, D. A., and Wright, P. E. (1989) Three-dimensional solution structure of a single zinc finger DNA-binding domain. Science 245, 635–637

